# Determining critical water potentials for creeping bentgrass seedling root elongation when exposed to PEG induced dehydration

**DOI:** 10.64898/2026.05.26.727908

**Authors:** Dominic Petrella, Maranda Morrow, Edward Nangle, Florence Breuillin-Sessoms

## Abstract

Creeping bentgrass (*Agrostis stolonifera*) is a turfgrass species established on golf course surfaces but is criticized for high irrigation requirements. While genetic variation for water deficit stress tolerance exists between cultivars, the lack of defined critical soil water potential thresholds (Soil Ψ_crit_) for this species complicates precise irrigation strategies and benchmarks for plant breeding. This study utilized a polyethylene glycol (PEG) infused agar-based system to simulate water potential reductions and determine the water potential threshold (Ψ_crit_) for seedling root elongation. Creeping bentgrass cv ‘Pure distinction’ seedlings were subjected to six water potentials (Ψ) ranging from -0.36 MPa (no PEG applied) to -1.72 MPa. Daily digital imaging was used to measure root elongation over 5 days. Results across two experiments demonstrated that creeping bentgrass seedlings are highly sensitive to mild reductions in Ψ. A reduction to -0.61 MPa significantly decreased root length and growth rates by over 50% compared to the control. Regression models predicted that a Ψ_crit_ of approximately -0.45 MPa reduced daily root growth by 25%, while upwards Ψ of -1.0 MPa resulted in a 75% reduction of seedlings root growth. Furthermore, seedlings exposed to the lowest water potentials were predicted to require an additional 30 to 46 days to achieve the same root length as control plants. These findings establish specific Ψ_crit_ benchmarks for water deficit stress tolerance using a PEG-based system to induce dehydration. These methods can be used in breeding programs, and will help develop more accurate experiments examining the mechanisms of water deficit stress tolerance.

## 1. Introduction

Turfgrasses are used worldwide for sports and recreational surfaces, for lawns, roadsides, cemeteries, and for many other purposes. While these species have been cited to provide many ecosystem and functional services for society, they are often criticized for requiring too much supplemental irrigation (Braun et al., 2022, 2024; Monteiro, 2017). This is especially true for turfgrasses used on golf courses. In cool-season climates, creeping bentgrass (*Agrostis stolonifera*) is the commonly established C_3_ species across different golf courses areas including putting greens, fairways, and tees. Creeping bentgrass can be maintained at low mowing heights (< 2.5 mm), withstands foot and vehicular traffic, and is relatively tolerant to temperature stresses (DaCosta & Huang, 2006; Young et al., 2015). However, the shallow root system and high evapotranspiration rates of creeping bentgrass lead to the need for frequent irrigation to prevent water deficit stress (Huang et al., 2014; Romero & Dukes, 2016). Even though creeping bentgrass is well suited for golf course surfaces, the amount of water this species requires is a growing concern, especially in areas where water is limited. Creeping bentgrass has been shown to have evapotranspiration rates as low as 3 mm d^-1^ but above 10 mm d^-1^ in summer months (Salaiz et al., 1991; Huang and DaCosta, 2006; Braun et al., 2022;). Furthermore, this species has been shown to exhibit genetic variation for tolerance to water deficit stress in field and controlled environment research, with the cultivars such as ‘Pure Distinction’ and ‘Penn-A4’ exhibiting greater tolerance compared to cultivars such as ‘Penncross’ and ‘L-93’ (McCann & Huang, 2008; Xu & Huang, 2010; Krishnan & Merewitz, 2022). Experiments examining creeping bentgrass tolerance have typically withheld water to simulate water deficit stress in both controlled environment and field experiments (Breuillin-Sessoms, et al., 2025; Breuillin-Sessoms, et al., 2025; Itam et al., 2024; Katuwal et al., 2022; Krishnan & Merewitz, 2022; Mengistu et al., 2025; Noor et al., 2024; Weldt et al., 2025; Yu et al., 2023; Zhao et al., 2026). In some instances, polyethene glycol (PEG) containing water has been irrigated onto creeping bentgrass to induce osmotic stress (Xu & Huang, 2010). Overall, these methods can make it more difficult to repeat experiments and examine genetic gain in breeding populations or new cultivars due to varying rates in the soil drying and soil types, leading to unpredictable timing of the onset of physiological stress.

While a range of water deficit stress tolerance mechanisms have been proposed for creeping bentgrass (Huang et al., 2014), to our knowledge, there is no defined critical soil moisture or critical soil water potential (Soil ⍰_crit_) for this species. A critical soil water potential is a threshold that when reached, the given plant will show signs of stress (Aronson et al., 1987). Defining ⍰_crit_ will allow for improved cultivar development by better benchmarking water deficit stress, improving testing throughput and irrigation precision while reducing water input.

For this research, a Petri plate-based method utilizing PEG infused agar was used to accurately simulate different water potentials. Creeping bentgrass seedlings were subjected to simulated water deficit stress in this PEG system to determine ⍰_crit_ using primary root elongation as a response metric. It was hypothesized that the varying PEG concentrations would elicit a direct response from the seedlings.

## 2. Materials and Methods

### 2.1 PEG infused agar-based media preparation

Polyethylene glycol (PEG) infused Murashige and Skoog (MS) agar-based media was prepared using previous published methods (Verslues et al., 2006) in 100 × 100 mm Petri plates. The media contained ½ strength MS basal salts (Caisson Labs, UT, USA) along with 2.56 mM 2-(N-morpholino)ethanesulfonic acid (MES) buffer, 0.75% Phytoblend agar (Caisson Labs, UT, USA), and a final pH of 5.7. Ultra-pure water (18.2 MΩ cm^-1^; Millipore Direct-Q 3 UV-R) was used for all steps including PEG solubilization. A total of 20 mL of liquid media was poured in Petri plates and solidified overnight within a biosafety cabinet (Labconco, Purifier Class II). Following this, 30 mL of a PEG solution was poured overtop the solidified gel. A total of 6 concentrations of PEG were applied, 0, 250, 400, 550, and 700 g L^-^1, using 8,000 M.W. PEG. Once poured onto the gelled agar within the Petri plates, the plates were sealed with parafilm and left within a biosafety cabinet for 15 hours to infuse. Following infusion, the remaining liquid PEG solution on top of the agar was discarded, and the Petri plates were used for testing.

### 2.1 Water potential of PEG infused agar-based media

The water potential of the PEG infused gels was evaluated using a dewpoint potentiometer (WP4C, Meter Group, Inc.) using five random Petri plate replicates of each PEG concentration. Data were analyzed using ANOVA with PEG concentration as a fixed effect and replicate as a random effect. Means were separated using Tukey’s HSD. Data were analyzed using JMP v18.0.

### 2.3 Creeping bentgrass PEG experiments

Creeping bentgrass cultivar ‘Pure Distinction’ seeds were surface sterilized using a previous method described in Petrella et al., 2018. Surface sterilized seeds were stratified at 10°C for two weeks within Petri plates on sterilized and moistened filter paper. Stratified Seeds were then placed on ½ strength MS agar (same as above) within 100 × 100 mm Petri plates and germinated vertically within a growth chamber (22°C constant temperature, 12 hr. photoperiod, 60-80 μmolm^-2^s^-1^). Four days after placement, rooted seedlings were transferred to sterile PEG-infused agar plates for treatment. PEG treatments consisted of 0, 250, 400, 550, and 700 g L^-1^ of PEG. For each PEG treatment, a total of 15 Petri plate replicates were used in a randomized complete block design, and 10 seedlings were placed on each Petri plate. Root elongation was measured daily over 5 days starting at day 0 (the day of transferring) by scanning plates with a flatbed scanner and measuring root length in ImageJ software (Schneider et al., 2012). The experiment was repeated twice, Experiment 1 and Experiment 2.

Each Petri plate was considered an independent experimental unit. Average root length was calculated for each plate, which was used for all downstream analyses. Average daily root length data were used to calculate 1) the average daily root growth rate, 2) the number of additional days required to elongate similarly to control plants growth, 3) the ⍰_crit_ for root elongation. Experiment 1 and experiment 2 were analyzed independently. All data were analyzed and figures produced in JMP v18.0 (Statistical Discovery LLC, USA).

#### 2.3.1. Daily root growth rate

For each PEG treatment, linear regression analysis were conducted to determine the daily root growth rate (calculated as the slope of the regression line). For this analysis, the water potential was estimated as a function of the PEG treatment. The fit curve platform with Petri plate placed within the “group” variable (i.e. random effect) and water potential placed within the “by” variable (i.e. fixed effect) was used. Calculated daily root growth rates were compared amongst treatments using a mixed model ANOVA with estimated water potential as a fixed effect and Petri plate as a random effect. Means were compared using Tukey’s HSD (Fig. 2 and 3).

#### 2.3.2. Additional days required for PEG treated roots to elongate similarly to control lengths

The slopes and intercepts of the linear regressions were used to calculate the number of additional days for roots grown in PEG infused agar media to reach 50, 75, and 100% of control (0 g PEG L^-1^) root length at the final time point. Final root length values of 1.92 cm and 1.80 cm were used for Experiment 1 and 2 respectively (as described in Table 1). For this analysis the water potential was estimated as a function of the PEG treatment. The number of days for roots to elongate were compared using a mixed model ANOVA with the estimated water potential as a fixed effect and Petri plate as a random effect. Means were compared using Tukey’s HSD.

**Table 1.**
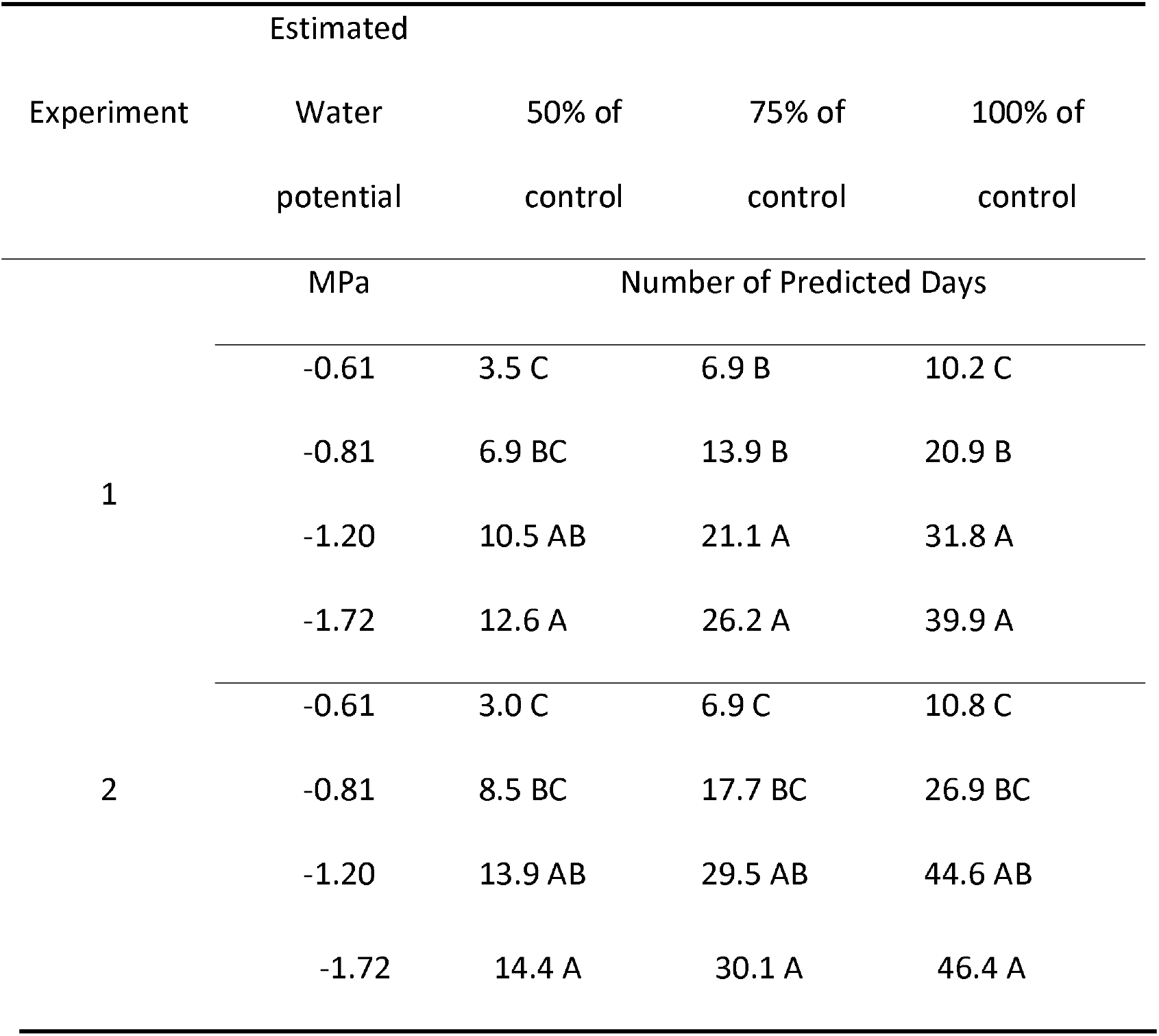
The predicted number of days for roots to elongate to 50, 75, and 100% of control roots (-0.36 MPa, no PEG added) at day 5 of treatment for Experiment 1 and Experiment 2.

#### 2.2.3. Critical water potential for root elongation

A three-point exponential regression was performed to determine ⍰_crit_ that reduced daily root growth by 25, 50, and 75%. The Fit Curve platform was used, and no data were assigned to group or by variables. Critical water potentials corresponding to 25, 50, and 75% reductions in growth were estimated using the custom inverse prediction function with 95% confidence intervals. Inverse predictions were constrained within the minimum and maximum water potentials tested, and no extrapolation was performed. The asymptote, scale, and growth rate parameters were used to define a reference 100% growth rate for the inverse predictions, which was 0.258 cm d^−1^ and 0.289 cm d^−1^ for Experiments 1 and 2, respectively (Sup. Fig. 1).

## 3. Results

### 3.1. Measure of Water potential in PEG concentration test

Petri plates with no PEG infused (0 g L^-1^) possessed an average water potential of -0.36 MPA. The average water potentials ranged from -0.36, -0.61, -0.81, -1.20, and -1.72 MPa for the different PEG treatments tested (Fig. 1). Statistical analysis confirmed that the measured water potentials in the different PEG infused treatments were significantly different at *p* < 0.0001.

**Figure 1.**
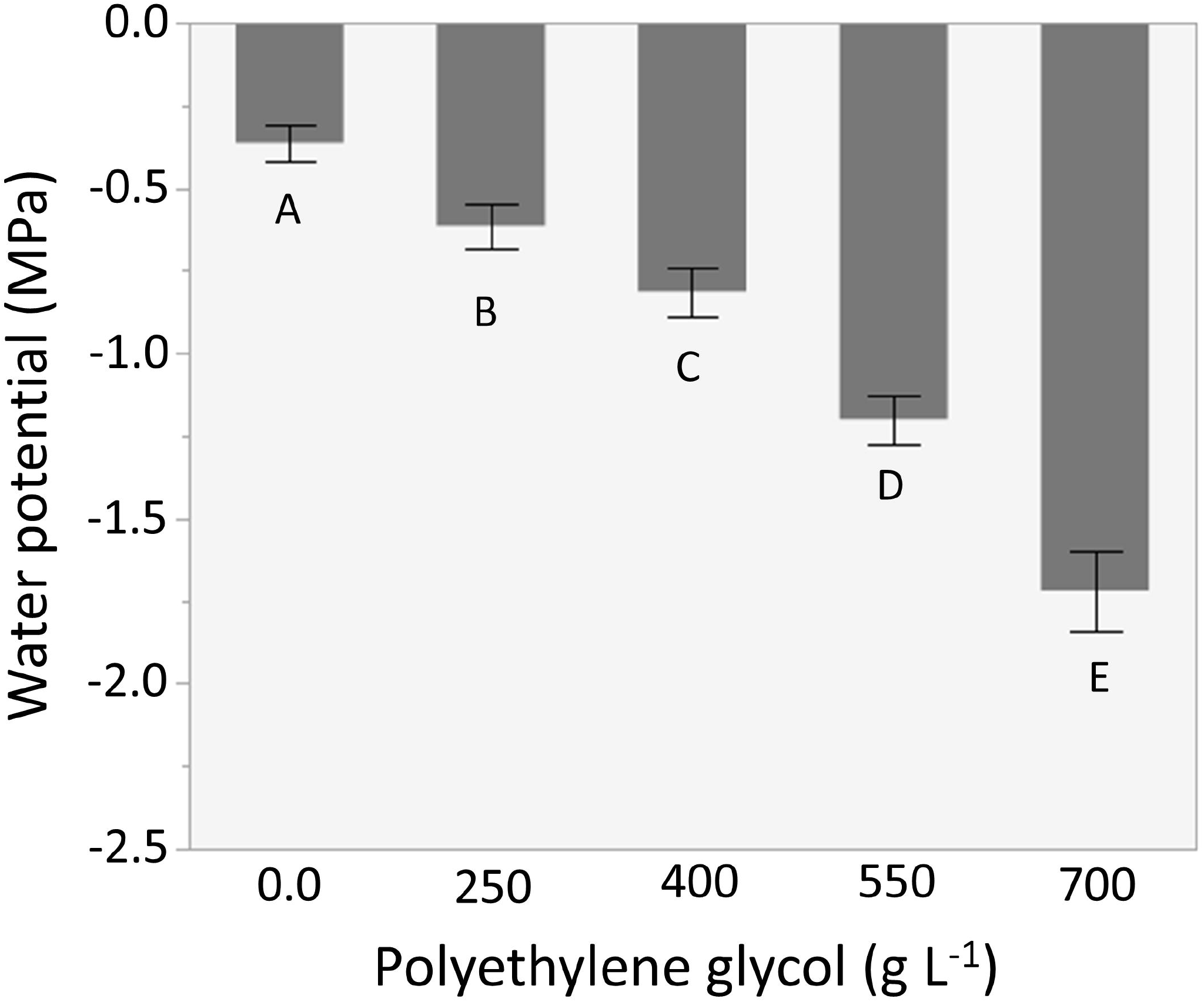
The effect of polyethylene glycol (PEG) added to overlay solutions on the final water potential (MPa) of agar gel within Petri plates. Average water potentials were –0.36, -0.61, - 0.81, -1.20, and –1.72 MPa for the different PEG treatments tested. Error bars represent 95% confidence intervals. Means were separated using Tukey’s HSD and means having different letters indicate statistical differences at the 0.05 probability level.

### 3.2. Daily root elongation

The root length of creeping bentgrass seedlings was significantly impacted by reduced water potential in both Experiment 1 and 2 (Fig. 2). A reduction of water potential to -0.61 MPa (red curve) significantly reduced root length compared to control (-0.36 MPa, black curve). Treatment with -0.81, -1.20, and –1.72 MPa severely reduced root lengths compared to control, but -1.2 and -1.72 MPa treatments were not significantly different from each other (Fig. 2) at p<0.0001.

**Figure 2.**
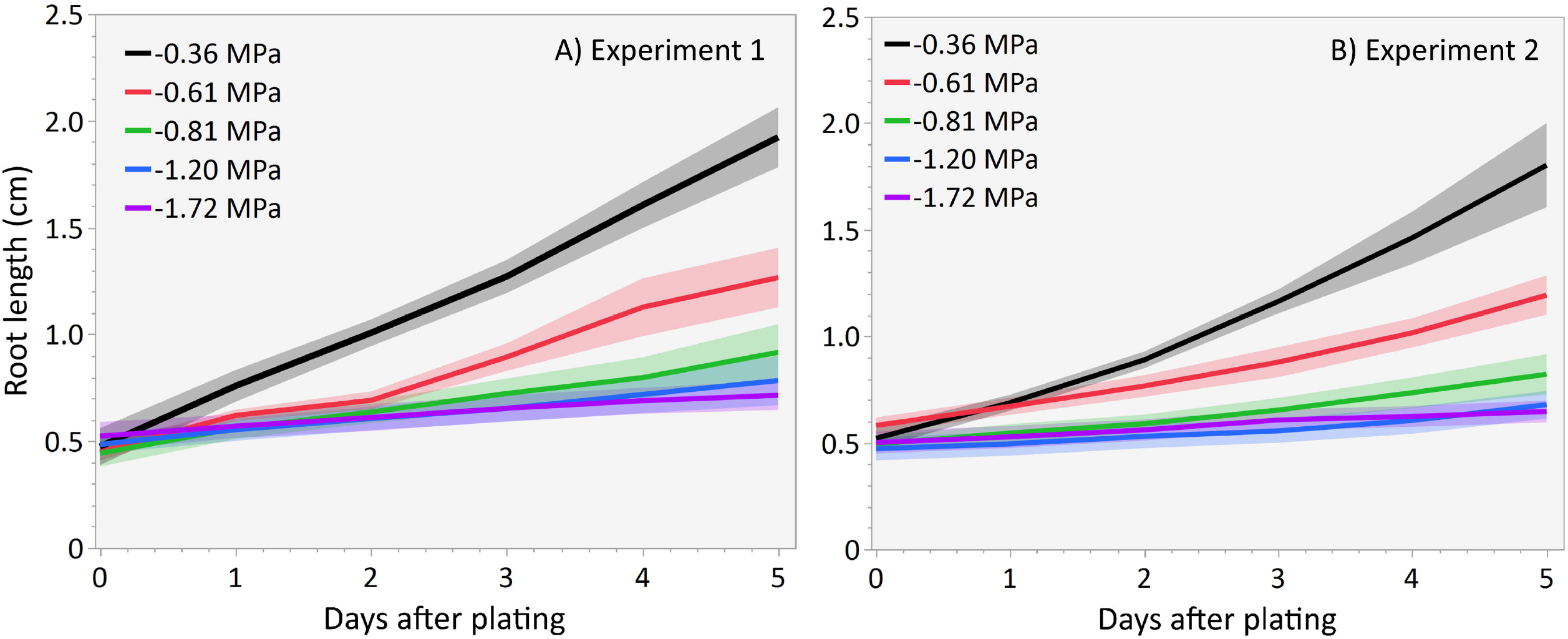
Average root elongation over 5 days in creeping bentgrass seedlings subjected to different water potential treatments for A) Experiment 1 and B) Experiment 2. Shaded areas represent the 95% confidence intervals for each treatment. Seedlings are 4 days old at placement in PEG infused Petri plates and 9 days old at the end of the measurement.

Calculated daily root length data for each water potential treatment was fitted into linear regression to determine the daily root growth rate. For these linear regressions, the coefficients of determination (R^2^) were higher than 0.94 in Experiment 1 and higher than 0.95 in Experiment 2. Water potential reductions significantly affected the calculated daily root growth rates for all treatments in which PEG was infused (*p* < 0.0001), with the mildest water potential reduction of -0.61 MPa already reducing the calculated growth rate by half or more (Fig. 3).

**Figure 3.**
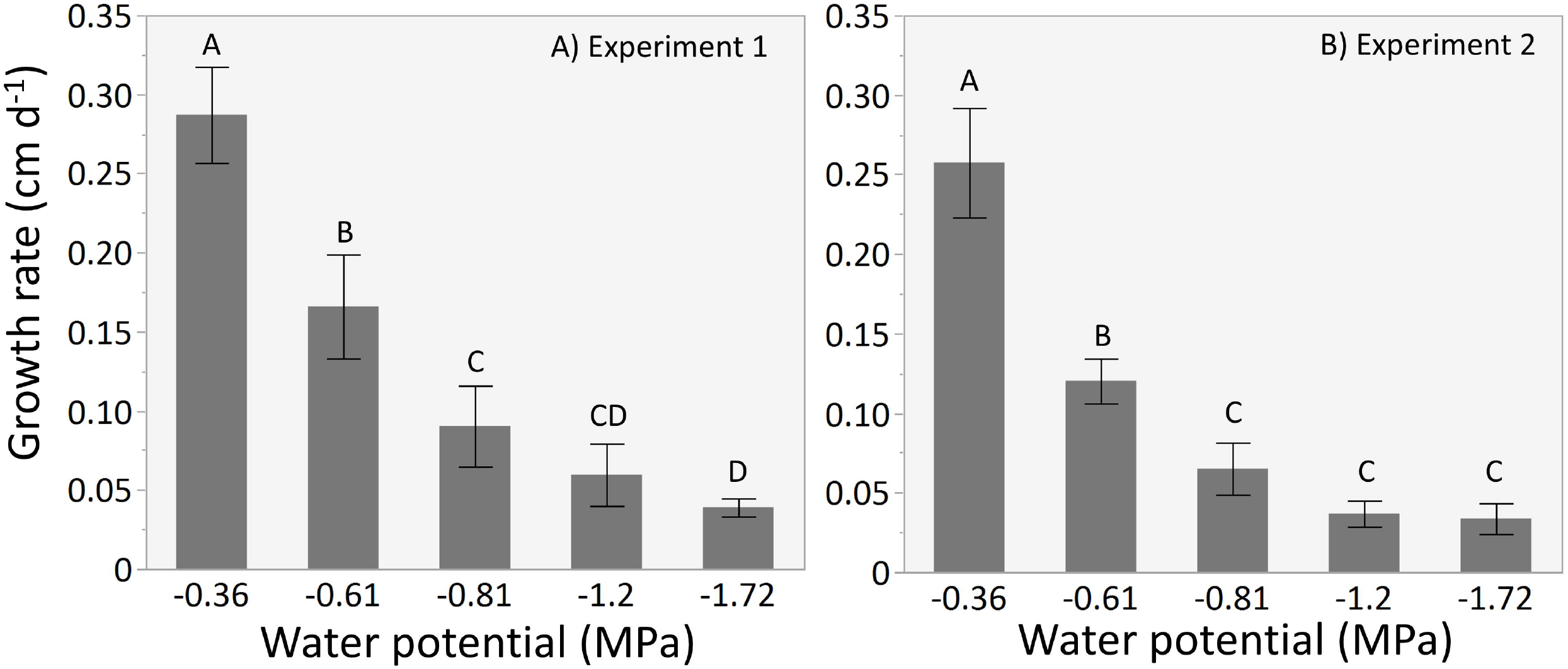
Calculated daily root growth rate (cm d-1) of roots as effected by water potential (MPa) for A) Experiment 1 and B) Experiment 2. Error bars represent 95% confidence intervals. Means were separated using Tukey’s HSD and means having different letters indicate statistical differences at the 0.05 probability level.

### 3.3. Determination of the additional number of days required for PEG treated roots to elongate similar to control

Creeping bentgrass seedlings with increasing concentration of PEG were predicted to require an additional 3, 6, and 10 days for roots to elongate to 50, 75, and 100% of control roots final length in both Experiment 1 and 2 (Table 1; (*p* < 0.0001)). Creeping bentgrass seedlings subjected to lower estimated water potentials showed a continuous increase of days to reach the 50, 75 and 100% of the final root growth in the control treatments (Table 1).

### 3.4. Determination of ⍰_crit_

The determination of the critical water potential (⍰_*cri*_) showed that creeping bentgrass seedlings were very sensitive to moderately low water potential. Critical water potentials of approximately -0.45 MPa were predicted to reduce daily root growth by 25% for Experiments 1 and 2, while critical water potential ∼-0.60 MPa reduce daily root growth by 50%, and critical water potential near -1.0 MPa was predicted to reduce daily root growth by 75% (Table 2).

**Table 2.** The predicted ⍰_crit_ (MPa) to reduce the daily root growth rates by 25, 50, and 75% for Experiment 1 and Experiment 2. Predicted water potential data are presented as 95% confidence intervals.

| Experiment | Relative reduction in | Critical Water |
| --- | --- | --- |
|  | daily growth rate | potential |
|  | % | MPa |
| 1 | 25 | -0.47 ± 0.03 |
| | 50 | -0.65 ± 0.05 |
| | 75 | -1.01 ± 0.12 |
| 2 | 25 | -0.45 ± 0.02 |
| | 50 | -0.57 ± 0.04 |
| | 75 | -0.84 ± 0.08 |

## 4. Discussion

This research showcase the responses of creeping bentgrass root growth to water potential reductions using increasing concentrations of PEG in a sterile Petri plate-based system. Considerable research has examined potential mechanisms that confer creeping bentgrass tolerance water deficit stress, but this study is one of few that determined critical water potential thresholds for this turfgrass species.

The water potential measured in agar gels that were infused with different concentrations of PEG were similar to what was shown in the previously described methods (Verslues et al., 2006) demonstrating the reliability and repeatability of the PEG infused method. Previous research has examined turfgrass and creeping bentgrass tolerance to water deficit stress in soil by various strategies of deficit irrigation and ET replacement (DaCosta & Huang, 2006; Sass & Horgan, 2006). These methods rely on measuring soil volumetric water content to correlate with plant stress, but ⍰ widely varies across different soil types. Different soils with the same water content will not have the same ⍰. In fact, the soil texture, structure, and organic matter will impose a strong influence on the soil water potential. Unfortunately, ⍰ is rarely measured directly or indirectly using water-retention curves (Novick et al., 2022; Osmolovskaya et al., 2018). The PEG infused method employed in this research is crucial for experiments or breeding where repeatability of water deficit stress is required. This is especially important for examining trait improvement in breeding populations where the stability of testing conditions is absolutely necessary to evaluate genetic gain and heritability.

Results from this study showed that creeping bentgrass seedlings are very sensitive to even small reductions in water potential with 25% reductions in growth predicted at -0.45 MPa. Interestingly, research has shown that a moderate reduction in water potential (-0.51 MPa), using PEG, stimulated root growth in *Arabidopsis* seedings van der Weele et al. (2000). However, that work had treatments where ⍰ was closer to zero, something this study did not have. Still, the high level of sensitivity observed in creeping bentgrass was also observed in mature *Arabidopsis* plants treated within a PEG infused agar. In this experimental set-up, -0.20 MPa was shown to lead to changes in plant growth (Frolov et al., 2017). Results from this previous investigation also demonstrate that this PEG infused system works on mature plants and not just seedlings. Further research with older creeping bentgrass plants (up to the tillering phase) should be conducted.

Previous greenhouse-based research evaluated the reduction of soil water potential defined on turf quality, relative evapotranspiration, leaf growth and relative leaf water potential (Aronson et al., 1987). The authors observed that a measured soil ⍰ of -80 kPa (-0.08 MPa) for Kentucky bluegrass (*Poa pratensis*) and perennial ryegrass (*Lolium perenne*), and -400 kPa (-0.40 MPa) or lower for Chewings fescue (*Festuca rubra* ssp. *Commutata*) and hard fescue (*F. brevipilia*) was necessary to reduce acceptable turfgrass quality (Aronson et al., 1987). It seems that in these experiments, the cultivars of Kentucky bluegrass and perennial ryegrass were more susceptible to the imposed drought stress that the creeping bentgrass cultivar in PEG induced water deficit. However, this previous study was run on silt loam soil with tensiometers placed at 10 cm to monitor soil water potential and may not capture the soil water potential many roots are growing; furthermore, breeding programs on these two species have tremendously increased their drought tolerance/resistance making comparison difficult to interpret.

In germination experiments, Goatley et al., (2017), found that a ⍰_crit_ reduced radicle length by 50% to average -0.78 MPa across a range of species. While there is little data in creeping bentgrass to compare the ⍰_crit_ that was determined from this research, water potential-root elongation growth curves in Maize show similar sensitivity to creeping bentgrass and ⍰_crit_ between -0.20 to -0.80 MPa (Sharp et al., 1988). Modeling research from Fu et al. (2021) also predicted mean ⍰_crit_ from diverse ecosystems that are in the range of what was also measured here, -0.71 MPa averaged across regions. As summer drought stress becomes more frequent and increasingly difficult to predict, PEG-induced water deficit systems offer a valuable tool for screening turfgrass germplasm and developing cultivars capable of growing under low water potential conditions. However, because PEG-induced stress differs fundamentally from soil-based drought stress, it cannot fully replicate the complexity of field environments. Therefore, field experiments remain essential to validate plant performance, ensuring that selected genotypes can tolerate and survive under real-world deficit irrigation conditions.

## 5. Conclusions

In summary, the findings from this research showed that creeping bentgrass seedling roots are very sensitive to small reductions in ⍰, with ⍰’s of ∼-0.60 reducing root growth by 50%. These data provide specific benchmarks and ⍰’s that are detrimental to creeping bentgrass and lay a foundation for future research. The PEG infusion system used here to simulate reductions in ⍰ worked well and could be reliably used for further research examining specific responses to low ⍰, helping to advance the development of creeping bentgrass cultivars with increased tolerance to water deficit stress.

**Credit**: **Dominic Petrella**: conceptualization; data curation; formal analysis; funding acquisition; investigation; methodology; project administration; supervision; validation; visualization; writing—original draft; and writing—review & editing. **Maranda Morrow**: data curation; analysis; investigation; methodology; writing—review & editing. **Edward Nangle**: funding acquisition; project administration; supervision; writing—original draft; writing—review & editing. Florence **Breuillin-Sessoms**: conceptualization; data curation; formal analysis; funding acquisition; investigation; methodology; project administration; supervision; validation; visualization; writing—original draft; and writing—review & editing.

## Acknowledgments

We thank FMC corporation for their supporting funding (AWD-114703).

## Conflict of interest

The authors declare no conflicts of interest.

---

Supplemental Figure 1. Three-point exponential regressions for the mean daily root growth rate (cm d-1, slope) and water potential (MPa) for A) Experiment 1 and B) Experiment 2. Equations for the regressions and root growth rate (GR) are inset into each panel.

## Notes

### Competing Interest Statement

The authors have declared no competing interest.

