## Supplementary figures and images for "Determining critical water potentials for creeping bentgrass seedling root elongation when exposed to PEG induced dehydration"

### Supplemental Figure 1

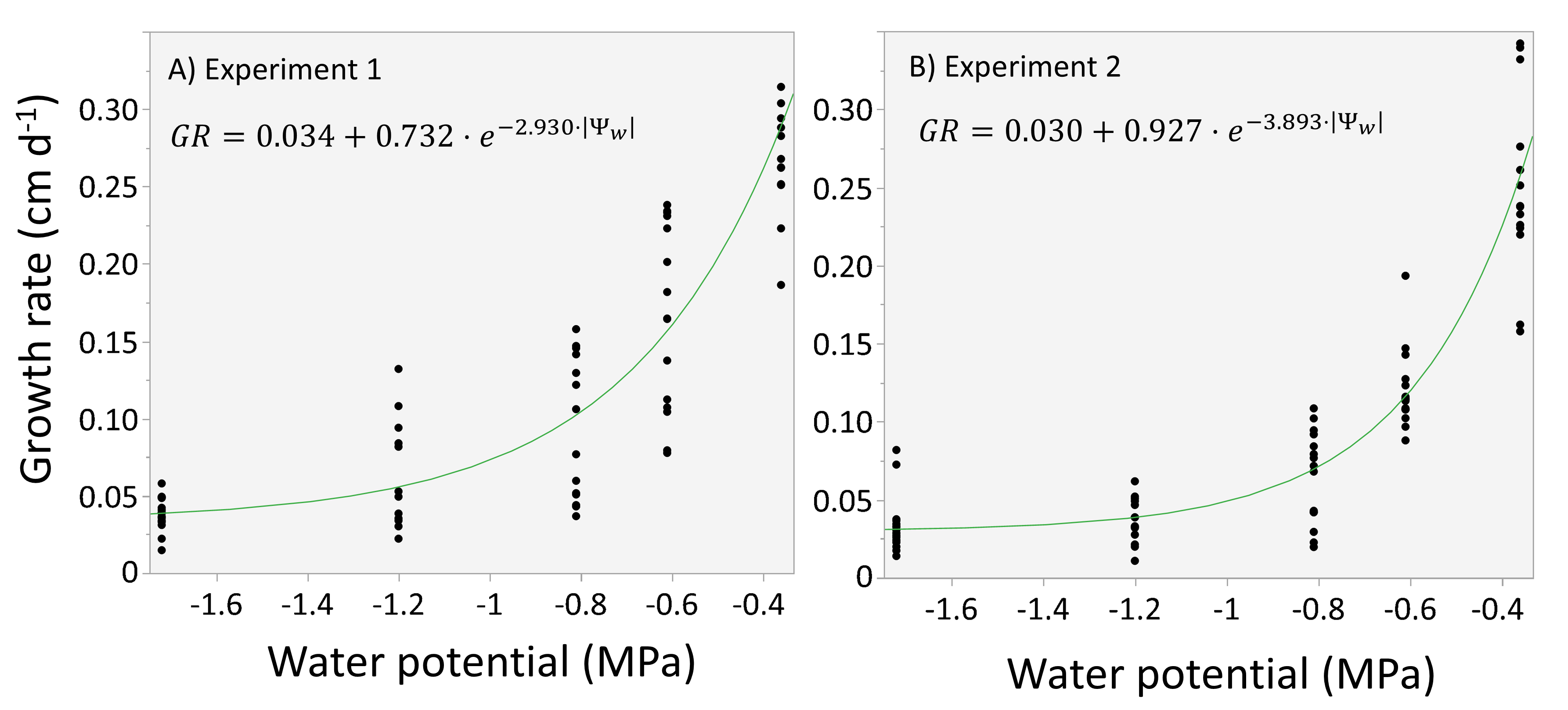
